# A collaborative submission model for building high-quality data resources at scale through partnership: lessons from single-cell genomics

**DOI:** 10.64898/2026.06.02.729387

**Authors:** Jason A. Hilton, Jim Chaffer, Jennifer Chien, Idan Gabdank, Brian Mott, Erica Rutherford, Corinn Small, Jennifer Zamanian, Brian Aevermann, J. Michael Cherry, Teri E. Klein

**Affiliations:** Department of Biomedical Data Science, Stanford University, Stanford, California, USA; Biohub, Redwood City, California, USA; Department of Genetics (emeritus), Stanford University, Stanford, California, USA; Department of Genetics, Stanford University, Stanford, California, USA; Department of Medicine, Stanford University, Stanford, California, USA

## Abstract

Community data resources that aggregate datasets across studies are critical infrastructure for modern biomedical research, accelerating discovery, and rapidly-advancing Artificial Intelligence (AI) models place ever-greater demand on large, well-described data corpora. However, building these resources involves a fundamental tension: comprehensive metadata is imperative for reuse, yet capturing this information systematically requires substantial effort that limits growth. We detail how CZ CELLxGENE Discover has used a collaborative submission model, whereby data contributors partner with dedicated resource curators, to become a large, widely used single-cell genomics resource for diverse analytical applications including AI model development and integrative analysis. This partnership leverages contributors’ intimate study knowledge and curators’ focus on data reuse and expertise in standardization to improve data quality, metadata accuracy, and richness. This is achieved by motivating researcher participation through tangible benefits while minimizing submission burden. We contrast this collaborative model with contributor-driven and resource-driven approaches, highlighting tradeoffs in scalability, consistency across submissions, and quality assurance. The principles and practices we describe provide a framework for building high-quality community data resources across diverse biological data types.

## Introduction

Public data resources are essential for maximizing the scientific value of research data: a centralized access point makes data from many studies findable and accessible, and shared standards make it interoperable and reusable. These FAIR (Findable, Accessible, Interoperable and Reusable) data principles [1] ultimately extend the lifecycle of data that might otherwise have little or no reuse after initial generation and analysis. Community resources can support hypothesis generation, supplement and validate newly generated data, and generate new insights through reanalysis alone [2]. In single-cell genomics, such resources let researchers contextualize findings across cell types, states, organisms, tissues, and conditions, and increasingly supply the substrate for computational and AI methods that consume data at scale.

However, data resources can struggle with a fundamental tension between quality and scale. Comprehensive, accurate, and standardized contextual information about experimental design, sample provenance, and data processing is critical for proper interpretation and reuse, yet capturing this information systematically across diverse studies requires substantial effort, creating a bottleneck as resources attempt to grow. Conversely, prioritizing rapid growth and scale often comes at the expense of thorough curation, leading to inconsistent annotations, missing contextual details, and reduced utility for cross-study analyses. Without sufficient metadata depth and quality, datasets become difficult to confidently integrate, compare, or reuse. This challenge is particularly acute in the single-cell genomics field, where experimental heterogeneity is continuously increasing with rapid technological advancements. Successfully building a broadly useful single-cell data resource therefore requires strategic choices about how data are aggregated in the resource.

One proven approach to balancing quality and scale is a consortium model, though its scope is inherently limited. Many of the most highly valuable and widely used resources for genomic data have been products of consortia, including those from ENCODE [3], HTAN [4], HuBMAP [5], and 4DN [6]. In the framework of a consortium, a data coordination center typically maintains a resource and works directly with data-generating groups starting as early as during experimental design, all operating under unified governance and shared scientific objectives. This organizational structure enables consortium portals to achieve both high data quality through standardization and effective scaling across the consortium’s research outputs. However, these resources are limited in scope to primarily compile data generated within the consortium itself, restricting their ability to aggregate the full breadth of publicly available data across the broader research community. The vast majority of research data is generated by independent research groups, rather than within consortium frameworks, and resources targeting the broader research community typically adopt one of two general strategies: contributor-driven submissions or resource-driven sourcing of public data.

Many resources establish a contributor-driven submission process that requires minimal, if any, involvement from resource personnel beyond the mechanics of data upload and basic format validation. These resources commonly have the primary goal of enabling researchers to share their data with the broader community; accordingly, they impose minimal standards and do not actively curate submissions to ensure accuracy, richness, or consistency with other datasets in the corpus. This approach can support the reuse of data from a given study but often makes cross-study analysis challenging. Examples span a spectrum of specialization: generalist repositories like Zenodo (zenodo.org) and Figshare (figshare.com) accept nearly any data type, domain-specific resources like the European Nucleotide Archive [7] and the Sequence Read Archive [8] cater to genomic data, and resources like the UCSC Cell Browser [9] and Single Cell Portal [10] focus on single-cell genomics data. Data contributors may prefer resources with the lowest submission burden, and generalist resources accept data from multiple domains, so a single submission yields a single access point for everything from a study. Thus, a contributor-driven approach lowers barriers to sharing and enables rapid corpus growth, but without rigorous curation and cross-study standardization, metadata quality and data formats vary widely, impeding integrative analyses even as emerging tools help contributors package their outputs more intentionally for reuse [11].

Resource-driven acquisition focuses on use cases requiring cross-study harmonization and sits at the other end of the spectrum, assembling a corpus from publicly available data without any involvement from the original data generators. Numerous genomic data resources [12–14] and single-cell genomics resources [15–23] employ this strategy. While this approach can produce well-curated, standardized corpora, it places a disproportionate burden on resource personnel, who are tasked with scouring literature and associated materials to identify suitable studies and reformat data products. This labor-intensive process severely limits scalability, as even well-resourced teams can comprehensively process only a small fraction of published datasets, yielding corpora that are internally consistent but represent an incomplete view of the research landscape.

AI-enabled software has the potential to counter some of these challenges by providing high-throughput tools to extract information, standardize metadata, and locate associated data products [19,24,25], but human curators and AI agents alike are limited by what information and data products have been shared openly. These constraints are especially pertinent in the single-cell genomics field where there is a lack of widely accepted data sharing best practices within the community [26]. For example, while the deposition of raw sequence reads is a common practice, and often a mandate, carried over from bulk genomics, this alone is insufficient for broad data reuse. Raw sequence data requires compute-expensive alignment, limiting accessibility to well-resourced groups, and raw sequence data from human samples may be under managed access for privacy reasons, further impeding reuse. A count matrix – an alignment product quantifying every reference-genome gene in every cell of the sample – is therefore vital to democratizing data access and enabling immediate reuse. This practical value of post-alignment products for reuse has recently been described across high-throughput sequencing assays more broadly [11]. Unfortunately, count matrices are not consistently shared with single-cell publications, so without contributor involvement, any resource providing standardized access to these matrices is either limited to studies that do share them or must invest substantial effort in realignment to generate them de novo. These challenges demonstrate why neither contributor-driven nor resource-driven approaches alone can efficiently build the high-quality, broadly useful data corpora that modern computational biology demands.

Here we describe a collaborative submission model that addresses the quality-scale tension without consortium-like organizational structure, building a valuable corpus from community submissions at CZ CELLxGENE Discover (hereafter CELLxGENE, pronounced cell-by-gene, cellxgene.cziscience.com) [27]. The resource is developed and maintained by engineers at Biohub, formerly the Chan Zuckerberg Initiative (CZI) and the “CZ” of its name, while we serve as the resource curation team. By centering submissions around count matrix data products, a minimal set of carefully-selected standardized metadata, and strong value propositions to motivate researchers, CELLxGENE builds a high-quality data corpus that serves diverse analytical use cases while maintaining scalability. Metadata richness, error correction, and cross-submission harmonization arise from contributors and curators iterating on each submission together, while sharing the workload keeps the process scalable. Grounded in our direct experience and supported by broader guidance on effective data portal design [28], the principles we describe apply to community resources serving research data of any type.

## Results

### Collaborative Submission Model

The collaborative submission model is based on active participation from both the contributors and a curation team dedicated to the resource, and an agreement from each on the final representation. The contributors’ first-hand knowledge of the research, combined with the curators’ resource expertise and focus on data reuse can ensure appropriate placement of each individual submission in the context of the whole corpus. A collaborative submission workflow can be arranged in an efficient, high-throughput manner that scales easily. No matter what specific tasks are assigned to each party, sharing the workload allows for a scalable submission process, resulting in an ever-growing data corpus even as submission requests increase.

CELLxGENE has successfully implemented a workflow around collaborative submissions in the absence of the funding structure provided by a consortium, and thus, is open to community submissions (**Figure 1**) [27]. CELLxGENE submissions begin with potential contributors contacting us with a request to submit their data. We confirm eligibility and communicate the submission requirements and process, followed by initial data preparation by the contributor. We then evaluate the prepared data, guided in part by a validator – a script that enforces the required data structure and metadata standards defined in the schema – and supplemental quality assurance tests (see **Code Availability**). From there, contributors and curators work together to clarify and refine the data as needed. Once agreed upon updates are finalized, the data can be released according to the contributor’s timeline.

**Figure 1.**
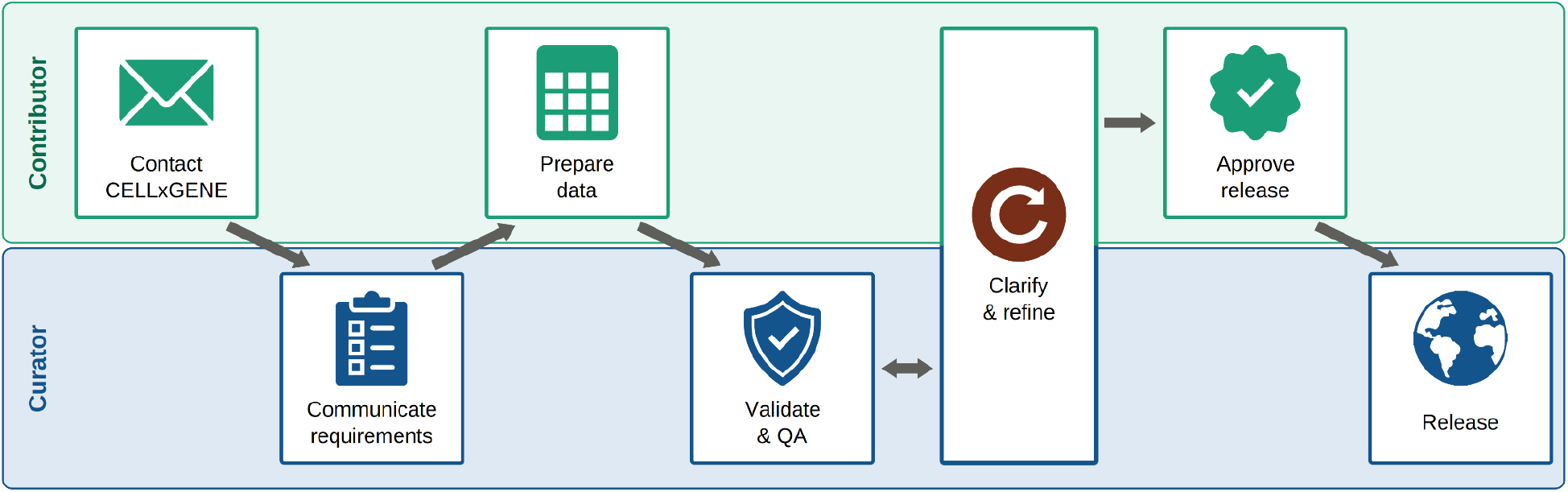
The CELLxGENE collaborative submission workflow. General steps and responsibilities are shown for contributor (upper lane) and curator (lower lane). Clarify & refine spans both lanes, indicating a step the contributor and curator work through together.

We maintain a structured submission workflow to ensure data consistency and quality, but we recognize that each contributor may have unique constraints or institutional requirements. Therefore, we implement a flexible approach that allows for case-by-case adaptations, when necessary, such as accommodating different methods for file transfer or providing hands-on support to help contributors map their metadata values to our required standards. Each submission serves as a learning opportunity to refine our processes, prompting simplification of steps that can reduce contributor burden and enhanced quality assurance checks based on identified edge cases. This commitment to continuous improvement ensures that the submission process remains as accessible and efficient as possible, lowering the barrier for future contributors and strengthening the foundation for a growing, high-quality data corpus.

### Evidence of Success

Since its launch in 2020, CELLxGENE has grown steadily to now encompass over 400 submissions, each released as a Collection, representing more than 170 million cells (**Figure 2a**). This sustained growth is possible because contributors, not curators, identify studies and draft submissions to meet our standards. Resource-driven approaches instead require personnel to review literature, search repositories for shared data products, and often learn only after extensive review that a study is ineligible. CELLxGENE curators still review manuscripts, protocols, and other documentation, but only to supplement what the contributor provides, and they help refine these drafts for datasets already known to be eligible.

**Figure 2.**
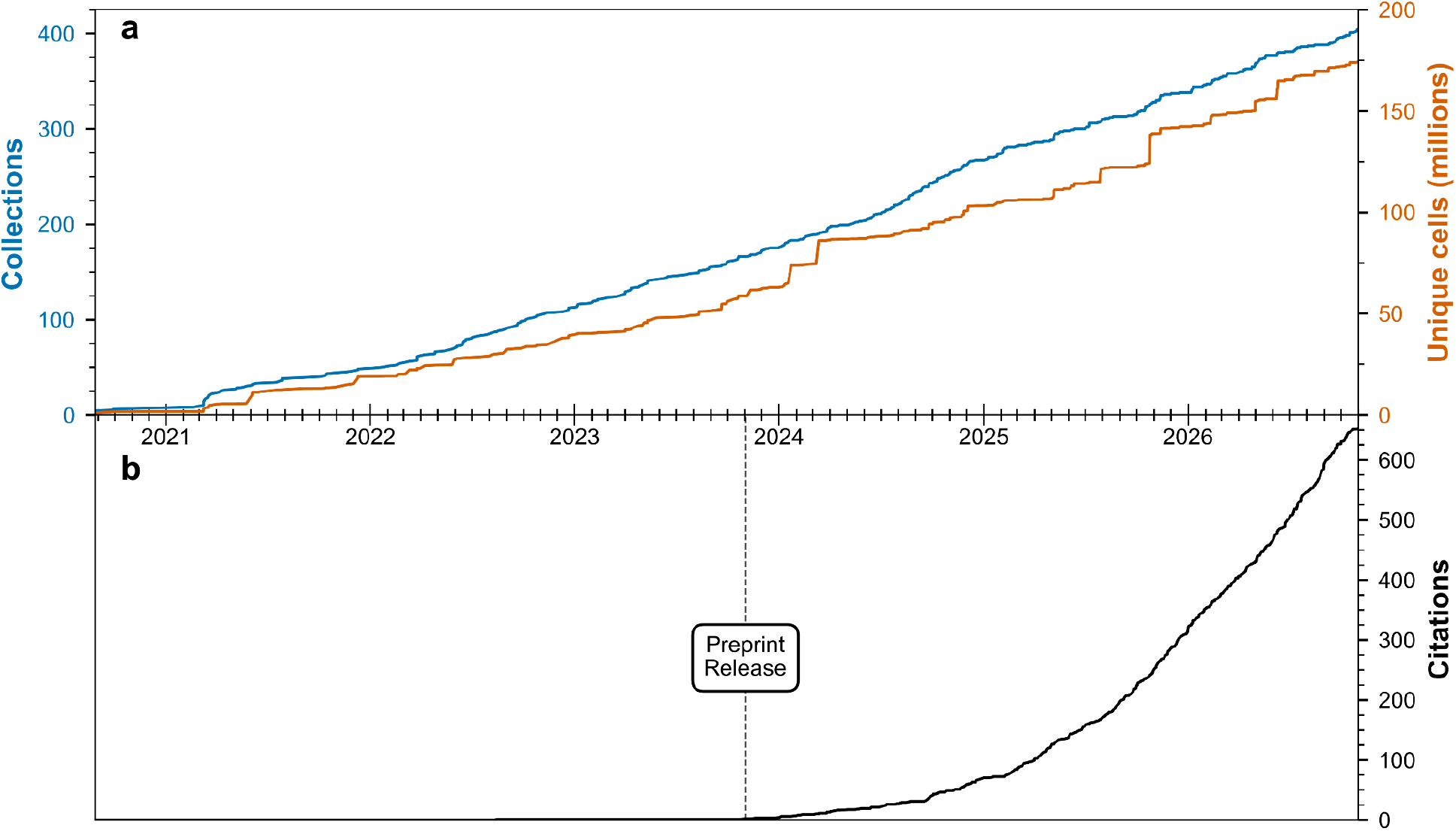
CELLxGENE corpus growth and usage. a) The cumulative counts of Collections, or submissions (blue) and unique cells (in millions; orange) released in CELLxGENE from August 2020 through October 2026, and b) the cumulative number of preprints and publications citing CELLxGENE from November 2023 through October 2026.

The CELLxGENE data corpus has maintained robust standardization as it has scaled, as exemplified by its heavy use by the community across diverse research applications (**Figure 2b**). The corpus is frequently used to train and test computational models and tools [29–39], compare the expression of specific genes across cell types and tissues [40–49], and enable multi-modal data integration studies [50,51]. This value is only achievable through the cross-study harmonization ensured by considerable attention given to each submission by curators. An enforceable schema is critical to this effort, but even well-defined standards will have gaps that allow for inconsistent interpretation, and a robust validator will not always prevent erroneous data from passing validation. Curators help contributors navigate the standards and offer clarification, when needed. For example, CELLxGENE requires submission of disease information, and this can be ambiguous for human studies – should all known participant diseases be documented, or only those meeting specific relevance criteria? Curators communicate a consistent definition of those criteria and ensure a standard application across studies.

Beyond guiding contributors through the standards, curators actively improve data quality by catching errors that would otherwise persist in the public corpus, supplementing the validator with a set of quality assurance tests run on every submission. While many of these tests are generalizable and can be run across submissions in a streamlined way, interpreting their results and recognizing the study-specific issues they surface requires a curator’s judgment rather than full automation. Some of the issues caught by these tests can directly impact the study’s analysis and interpretation, like samples inadvertently duplicated within an analysis or donors with misreported sex. In this way, the curators perform a peer review on the data itself. While peer review of publications is standard practice, peer review of the underlying data is not, but it is a standard part of every submission to CELLxGENE. Because curators are working together with the contributors on the submission, and only because of this partnership, the researchers can answer questions and resolve any issues so they can be corrected before making the data publicly available and analyses can be updated before publication, if needed. Contributors also catch previously overlooked errors when viewing their reformatted data in the CELLxGENE interface, a fresh representation that breaks the visual familiarity of their own working environment, where repeated exposure can obscure mistakes.

The iteration between curators and contributors during submission also commonly results in increased metadata richness. Curators are able to help contributors refine their metadata by directly conveying the benefits that downstream data consumers will gain by using more specific terms. This commonly arises when contributors of human data cannot share, or do not know, specific donor ages and thus submit “unknown,” which is valid but intended only when no information is available. In many of these cases, however, some information is known, whether it’s an age range or that all donors were at least of a certain age. Curators help identify the appropriate developmental stage terms that reflect the known information and will provide much more value to downstream users than “unknown”. Beyond selecting the most appropriate existing terms, curators can also identify when the ideal scenario is to expand or refine the standards themselves. When a contributor wishes to describe an experimental variable not yet seen in previously submitted studies, curators can identify a path to avoid having the contributor default to a less descriptive term. Curators commonly do this by submitting new term requests with the ontologies underlying the CELLxGENE standards. Without curator guidance, contributors may lack insight into what downstream users need and thus focus on meeting minimum validation requirements without recognizing opportunities to provide additional nuance and precision.

Curator guidance extends beyond metadata refinement to the data products themselves, ensuring they are prepared in ways that maximize reusability. For example, when a contributor provides a matrix filtered to the genes retained through their analysis workflow, curators explain the value of including the full, unfiltered gene set. The genes vital to the contributor’s analysis may not be the optimal gene set for a downstream user’s analysis, and by collecting unfiltered data, CELLxGENE enables diverse analytical approaches. In contrast, contributor-driven and resource-driven approaches are limited, short of realignment, to whatever gene set the original submission included, which may be heavily filtered for the initial study, leaving downstream users a narrower analytical foundation than the data could support.

Identifying overlapping data across the corpus, and thereby preventing inadvertent duplication in downstream analyses, requires participation from both contributors and curators. To do this, for each sample in every submission, the publication describing the original data generation must be recorded. Integrative studies typically detail which data were reused, but studies that combine newly generated with previously published data often make that reuse less clear in the metadata and sometimes in the manuscript itself, particularly when the earlier data came from the same research group. Contributors help curators identify these original studies, and curators compare them against every study already in the corpus. Only this collaboration lets CELLxGENE flag data appearing in multiple submissions and keep downstream users from unknowingly analyzing the same cells twice.

The value of these contributor-curator relationships extends beyond individual submissions. To maintain a harmonized corpus over time, not only is submission-to-submission consistency vital, but re-curation of previously submitted data may also be needed if any standards are added or adjusted. Resources should always strive to avoid breaking changes that invalidate existing data, but they are sometimes unavoidable when evolving based on user feedback or emerging use cases. CELLxGENE has had one notable update, a schema version that introduced requirements for donor identifiers and suspension type (schema v3.0.0, implemented November 2022; see **Code Availability**), which resulted in corpus-wide re-curation. In this instance, curators alone were able to extract the necessary information for many of the submissions, but the remaining datasets required input from the researcher. Remarkably, the response rate for these cases was 100%, and we were able to fully migrate the corpus to the updated schema version. Without the familiarity formed during submission, researchers would be far less likely to respond, and expanding requirements would likely have cost some data that could not meet the updated standards, leaving consumers with an inconsistent or incomplete corpus.

CELLxGENE measures success not just through the value provided to consumers of the data, but also through the value realized by contributors sharing their data in the resource. Most CELLxGENE submissions come from researchers wishing to share their data simultaneously with the publication as a way to emphasize their findings. However, as of November 2026, one third of submissions were released more than 200 days after their associated publication, showing that contributors see value in CELLxGENE independent of publication timing. Additionally, out of 305 contributors, 53 have returned to submit more datasets to CELLxGENE, a strong indication that the value gained outweighs the burden of submission. These metrics, together with frequent submission requests (more than 7 eligible per month over the last 12 months, plus requests for ineligible data), indicate that the tangible benefits of submission overcome common reluctance to share.

The value contributors gain, however, extends beyond the submission itself. Direct engagement with curators and the standards gives contributors firsthand exposure to data sharing best practices, such as using community ontologies and understanding which data products best support reuse, that they carry into future work, including submissions to resources with fewer or no standards. In this way, the collaborative submission model can contribute to a broader cultural shift toward more rigorous and meaningful data sharing.

CELLxGENE actively reinforces these practices structurally, grounding its standards in community ontologies and aligning them with the broader biomedical data ecosystem. Furthermore, several resources in the scFAIR initiative have adopted the CELLxGENE schema [52], creating cross-repository alignment and facilitating cross-resource analyses.

### Implementation Guidance

Understanding and addressing data sharing reluctance is vital to a collaborative submission model because success relies on motivating scientists. The prospect of increased citation and supporting downstream reuse can motivate contribution to a larger corpus, but a more powerful value proposition elevates the study at hand, as CELLxGENE and other single-cell resources do by turning static dimensionality-reduction figures (e.g. UMAP embeddings) into interactive versions [27]. In the single-cell community, many researchers create their own portal or app for such visualizations [53–56], but by submitting data once to CELLxGENE, researchers avoid the burden of installation, long-term maintenance, and continued data storage. This is possible because CELLxGENE represents contributors’ analyses as performed rather than reprocessing data through uniform pipelines. The data products in CELLxGENE directly correspond to those underlying the published findings, and researchers can confidently point to their submission as the authoritative representation of their study. Uniform processing pipelines are necessary for some cross-study analyses, but resources that reprocess data cannot offer this contributor value, since their data products deviate from the contributor’s own analysis. By offering tangible benefits that enhance the visibility and accessibility of contributors’ work, a resource can successfully motivate researchers to invest effort in data submission.

Beyond the data products themselves, contributor motivation also depends on flexibility in metadata representation. It is imperative that contributors are given the opportunity to describe their data in ways that they want, albeit so long as it does not jeopardize the desired cross-study standardization within the corpus. As we discuss below, CELLxGENE aims for minimal requirements and enforces those strictly. However, contributors may add any additional information in a format of their choosing, to group or label data as they prefer or to record details of samples, experimental model, and analysis not covered by required fields. The balance between standardization and flexibility ensures contributors can represent their work authentically, reinforcing their sense of ownership over the submission and their motivation to share.

CELLxGENE further supports contributors by letting submissions remain private while they and their colleagues review and iterate, and so peer reviewers can have access before public release [27]. Data can therefore be submitted pre-publication, released alongside the corresponding publication, and cited there by URL, which also increases awareness of the resource. This timeline alignment also satisfies sharing mandates for resources hosting data products required by journals or funders.

Even a motivated contributor can be deterred by an onerous submission process, so it is imperative to minimize submission burden, in part by limiting metadata requirements. Resources must balance the information desired by data consumers against what is readily available to contributors and the effort required to get that information into the standard format. Data consumers commonly request extensive standards, but every added requirement deters submissions, leaving those same consumers with a smaller corpus. Consumers will equally be ill-served by required fields that lack standardization. Where submission-to-submission variability makes it challenging to translate information into enforceable standards, resources must fall back on free-text fields, which inevitably results in the same concepts inconsistently represented across submissions and undermines cross-study harmonization. Downstream value therefore justifies a new requirement only when its consistency can be enforced across submissions.

CELLxGENE accordingly requires ten properties to capture information about the donor, sample, and experiment, and we generally introduce new requirements only when supporting new data types (see **Code Availability**) [27]. We prioritize information consumers need to identify studies of interest (e.g. species, anatomical region sampled, and disease) alongside variables known to impact measurements and enable cross-study comparison (e.g. suspension type, library preparation kit, cell type, and some donor metadata). The schema further requires stable gene identifiers so that features are directly comparable across submissions and the information required to create visualizations. A count matrix is required for RNA-seq submissions and a fragments file for ATAC-seq, centering each assay on the post-alignment but otherwise unprocessed product that can democratize access to analysis-ready data. Other information is valuable but resists consistent representation across submissions, and we exclude it from our requirements, prioritizing standardization over comprehensiveness. For example, processing metadata such as pipeline software and version and reference transcriptome would benefit many downstream users, but the high variability and customization of pipelines across studies makes enforceable standardization impractical. However, the CELLxGENE schema allows contributors to supplement the required fields with any optional fields, so that minimal requirements do not come at the cost of study-specific details. Those details, together with the link from each submission to its associated publication, mean consumers whose needs exceed our standards still benefit from centralized, consistently structured data as a starting point.

To make sound judgment calls about what to require, curators should be unencumbered by their own analytical interests in the corpus, free to weigh a variety of use cases rather than overfitting the resource to one type of analysis. That independence also protects the bandwidth to assist and troubleshoot each submission. The CELLxGENE curators are staff scientists whose work across projects converges on making research data more accessible. Because even well-defined standards leave room for interpretation, consistency depends on curators knowing what submissions others are managing. With an average tenure of 4.9 years, we have found stability and cohesion are essential to that.

Documentation precedes any contact with curators and often introduces a resource to the community, so it must be clear and concise. Complexity in documentation can deter users before they can offer the feedback that would expose weak points. CELLxGENE publishes the full schema in GitHub (see **Code Availability**), but keeps the entry point simple, providing a summarized outline of requirements (cellxgene.cziscience.com/docs/032 Contribute%20and%20Publish%20Data). Once contact is established, contributor input informs submission processes, but also informs the development team’s work on portal features for both contributors and data consumers. CELLxGENE draws on contributor relationships when designing features – explicitly through direct questioning, implicitly through understanding accumulated over prior interactions.

While insights gained through contributor interactions are invaluable, sustaining and growing the resource also requires proactive engagement with the broader community. CELLxGENE maintains multiple channels for engagement, including a Slack workspace and help-desk email for ongoing user support, where users can reach resource personnel, including the curation team. The curation team also regularly attends conferences to connect with both potential contributors and data consumers. This outreach reveals what value contributors seek, what metadata consumers need, and what new data types to support, while enabling ongoing troubleshooting and building trust within the research community. Critically, outreach allows curators to engage with researchers early in their research process, when data sharing can be integrated into their full workflow plan rather than addressed as an afterthought during manuscript submission. Early engagement increases the likelihood that researchers will both be aware of the resource and allocate sufficient time to meet submission requirements. By maintaining close connections with the community through outreach, a resource can ensure that the collaborative submission model remains responsive to the evolving needs of both data contributors and consumers.

### Challenges and Limitations

The most significant hurdle to building a resource through collaborative submissions is getting started. Without an established track record or sizable corpus, the resource faces a dual challenge: the community lacks awareness of its existence, and potential contributors may lack trust in its value and longevity. Researchers are more likely to invest effort in submission when they can see example datasets, recognize peers who have contributed, and observe that the resource has gained traction within their field. For newer communities or emerging data types, like the single-cell genomics field, this challenge is compounded by the absence of established data sharing standards and journal requirements. Without such external motivators, the resource must both tap into what researchers already value and actively shape new practices, demonstrating the value and feasibility of sharing data products (like count matrices) and metadata outside researchers’ typical workflows. This norm-setting role means early submissions require overcoming inertia around existing habits rather than simply meeting recognized requirements.

Therefore, it may be necessary to make some concessions on the submission process early on for the sake of establishing an initial data corpus. Standard CELLxGENE submission process involves the contributor preparing the data products with standards, but many early CELLxGENE submissions were initially prepared by the curators. In these cases, the researchers were only contacted once a draft submission was ready in an effort to minimize the burden of submission on the contributor. The visualization provided for each dataset submitted to CELLxGENE, even in draft form, can be an exciting motivational tool. From there, all that was asked of the contributors was to review the submission and possibly answer a few clarifying questions. More curator bandwidth was required to review possible studies for submission and prepare the draft submission, but we exchanged that effort for higher likelihood of the submission being completed to release. A resource must be careful not to compromise too much while trying to catalyze a data corpus, but some strategic sacrifices in early development can initiate a strong foundation for the future.

One specific strategy with these early CELLxGENE submissions was to prioritize preprints so that the submission would be ready in time for the resource URL to be shared with the publication, which brings much-needed awareness to the resource. We also targeted larger multi-lab projects such as consortia for early partnerships, both for their data volume and because a single partnership brings visibility and relationships with multiple research groups. The BRAIN Initiative [57] was a particularly pivotal partnership formed in the early days of CELLxGENE. Several large datasets from The BRAIN Initiative were ready for release within the first six months that CELLxGENE was accepting submissions, and the high-profile nature of the project and the specific studies helped bring critical awareness to the young resource.

Building on this foundation, CELLxGENE established partnerships with other major initiatives, each bringing new data, visibility, and credibility that are essential to attracting more contributors.

Leveraging existing community relationships and institutional credibility can significantly accelerate a new resource’s early growth and adoption. CELLxGENE benefited significantly from such groundwork, as the “CELLxGENE” name was already familiar in the single-cell community as a standalone software package for single-cell data visualization and annotation, now termed CELLxGENE Annotate [58]. Additionally, Biohub had fostered extensive connections through various grant programs, meeting sponsorships, and other community engagement activities. The trust cultivated through these relationships extended to the resource, providing the curation team with an initial list of potential contributors and access to user research that informed requirements development and identified motivations for data submission. This foundation substantially shortened the runway needed to build a robust data corpus.

Additional drawbacks of a collaborative submission approach include the need for continued support of a dedicated curation team. Our team has other data coordination duties, but we estimate that CELLxGENE work – submissions, user outreach, and schema expansion and validation with Biohub engineers – is equivalent to roughly 4 full-time staff scientists. As AI agents advance in capability, there will be opportunities to alleviate curator burden in areas like quality assurance test design and execution. However, AI agents remain limited by what has been published and shared, and lack the iterative feedback loop with contributors that drives metadata refinement, error correction, and process improvement.

The dependence on contributor engagement that makes the collaborative model so effective also introduces an inherent limitation in corpus composition. CELLxGENE occasionally has some bandwidth to solicit specific researchers to submit data, and the curation team can target specific conferences for outreach opportunities but the resource is ultimately dependent on submissions from motivated contributors rather than collecting any study of interest. While our community-driven submission approach has advantages, it means the resource’s composition is shaped more by contributor willingness than systematic curation priorities, such as ensuring inclusion of high-impact studies and balancing representation across different biological systems and study contexts. This is another area where resources may benefit from AI agents in identifying significant gaps in the corpus and published studies that might be solicited for submission to address those gaps.

## Discussion

Large, well-described bodies of published data, like CELLxGENE’s corpus, catalyze scientific discovery and are becoming increasingly valuable as computational tools rapidly scale in their ability to consume, analyze, and interpret data. Realizing that value, however, requires motivating researchers to share their data in a meaningful and intentional way, rather than simply depositing files to satisfy a requirement. At CELLxGENE, we overcome the common reluctance scientists have toward data sharing by implementing collaborative submissions that provide tangible value to individual contributors while simultaneously ensuring both the data products and standardized metadata needed for diverse cross-study analyses. We strongly encourage community resources to consider a similar collaborative submission approach as a means to build these data corpora – a need recognized in fields well beyond single-cell genomics, including bioimaging [59,60]. As AI agents advance, direct collaboration between researchers and curators remains essential for capturing the data products and contextual metadata that publications alone cannot provide, enabling a resource to achieve both cross-study standardization and efficient scaling while serving contributors’ data sharing needs with publication. With very intentional strategy and thoughtful design, a resource can return enough value to researchers and build sufficient trust within the community to motivate submissions. Each submission strengthens the corpus, and the corpus strengthens the case for the next, establishing a cycle that continuously expands the foundation upon which scientific discovery can be accelerated.

## Data Availability

CELLxGENE data are openly accessible: cellxgene.cziscience.com.

## Code Availability

The CELLxGENE schema and validator: github.com/chanzuckerberg/single-cell-curation.

Current CELLxGENE schema: chanzuckerberg.github.io/single-cell-curation/latest-schema.html.

All schema versions: github.com/chanzuckerberg/single-cell-curation/tree/main/schema.

The supplemental quality assurance tests developed and used by CELLxGENE curators: github.com/Lattice-Data/lattice-tools/blob/main/cellxgene_resources/curation_qa.ipynb.

The code to generate Figure 2 and the submission statistics cited in this paper: github.com/Lattice-Data/lattice-tools/tree/main/cellxgene_resources/collab_submissions_pub.

## Acknowledgments

We are grateful to Seth Strattan whose mentorship and guidance was instrumental in shaping how we approach community collaboration and data standardization. We also thank the many Biohub personnel who have contributed to designing, building, and maintaining the resource, establishing partnerships, and providing ongoing scientific direction, especially Brian Raymor. Lastly, we thank all the data generators and data consumers who have collaborated with us for their insights and commitment to open science.

## Author Contributions

JAH, BA, JMC, and TEK designed the research. JAH wrote the manuscript. All authors commented on the manuscript. JAH, JChaffer, JChien, IG, BM, ER, CS, and JZ executed the curator roles described.

